# CytoGLMM: Conditional Differential Analysis for Flow and Mass Cytometry Experiments

**DOI:** 10.1101/2020.12.09.417584

**Authors:** Christof Seiler, Anne-Maud Ferreira, Lisa M. Kronstad, Laura J. Simpson, Mathieu Le Gars, Elena Vendrame, Catherine A. Blish, Susan Holmes

## Abstract

**Background:** Flow and mass cytometry are important modern immunology tools for measuring expression levels of multiple proteins on single cells. The goal is to better understand the mechanisms of responses on a single cell basis by studying differential expression of proteins. We focus on cell-specific differential analysis and one fixed cell type. In contrast, most current methods learn cell types and perform differential analysis jointly. Our narrower field of application allows us to define a more specific statistical model with easier to control statistical guarantees.

**Results:** Differential analysis of marker expressions can be difficult due to marker correlations and inter-individual heterogeneity, particularly for studies of human immunology. We address these challenges with two multiple regression strategies: A bootstrapped generalized linear model and a generalized linear mixed model. On simulated datasets, we compare the robustness towards marker correlations and heterogeneity of both strategies. For paired experiments, we find that both strategies maintain the target false discovery rate under medium correlations and that mixed models are statistically more powerful under the correct model specification. For unpaired experiments, our results indicate that much larger patient sample sizes are required to detect differences. We illustrate the `CytoGLMM` *R* package and workflow for both strategies on a pregnancy dataset.

**Conclusions:** Our approach to find differential proteins in flow and mass cytometry data reduces biases arising from maker correlations and safeguards against false discoveries induced by patient heterogeneity.

## 1 Introduction

Flow (Saeys, Van Gassen, and Lambrecht 2016) and mass cytometry (Bendall et al. 2011) allow researchers to simultaneously assess expression patterns of a large number of proteins on individual cells, allowing deep interrogation of cellular responses. The goal of such studies is to better understand the mechanisms of responses on a single cell basis by defining protein expression patterns that are associated with a particular stimulus or experimental condition. Finding differentially expressed proteins can help identify how cells function across experimental conditions.

Statistical workflows to analyze data generated by flow and mass cytometry usually begin by clustering cells into both known and novel cell types. Many cluster algorithms are available (Lo et al. 2009; Finak et al. 2009; Qian et al. - Zare et al. - Aghaeepour et al. 2011; Qiu et al. 2011; Ge and Sealfon 2012; Shekhar et al. 2014; Becher et al. 2014; Naim et al. 2014; Meehan et al. 2014; Van Gassen et al. 2015; Sörensen et al. 2015; Levine et al. 2015; Chen et al. 2016; Samusik et al. 2016; Lee et al. 2017; Li et al. 2017; Theorell, Bryceson, and Theorell 2019; Abdelaal et al. 2019) and Weber and Robinson (2016) provide an informative benchmark comparison study of most of these algorithms. The cluster step is followed by a differential expression analysis between and within cell types. The most popular differential analysis tools are: Citrus (Bruggner et al. 2014), the `Bioconductor workflow` by Nowicka et al. (2017), `cydar` (Lun, Richard, and Marioni 2017), `CellCnn` (Arvaniti and Claassen 2017), and `diffcyt` (Weber et al. 2019).

We can classify differential analysis methods into marginal regression—analyses that focus on individual markers—and multiple regression—analyses that work on multiple markers simultaneously. The `Bioconductor workflow` by Nowicka et al. (2017), `cydar`, and `diffcyt` are marginal regression methods. The advantage of marginal regression approaches is that they allow for flexible experimental designs. The main disadvantage of this approach is the separate testing for differential expression for each protein—when studying a specific protein marker all the other markers are ignored. Therefore these methods are susceptible to biases induced by marker correlations.

`Citrus` and `CellCnn` are multiple regression methods. The advantage is that they can provide a conditional interpretation of the effect of a protein onto the outcome, and thus reduce the bias coming from marker correlations. The disadvantage is that `Citrus` summarizes protein expressions by taking the median for each cell type which can lead to a decrease in statistical power. The power decrease comes from the reduction in cell sample size from thousands of cells to one cell per sample. On the other hand, `cydar` uses a neural network for which it is currently unclear how to build confidence intervals, derive *p*-values, and control the number of falsely reported markers.

It is helpful to consider an example to further illustrate the differences between the marginal and the multiple regression method. Consider two intracellular proteins, *A* and *B*, that are part of the same signal transduction pathway. Assume that applying a stimulus to *A* activates *B*. Further assume that the stimulus does not directly activate *B*. If we performed separate differential analyses on protein *A* and *B*, we would observe differential expressions for both *A* and *B*, even though only *A* had been directly activated. In contrast, a multiple regression method would report *A* as differentially expressed given *B*, and *B* as not differentially expressed given *A*.

`CytoGLMM` implements multiple regression that accounts for marker correlations without the aforementioned limitations. The main difference between our method and current methods is that we focus on cell-specific differential analysis and one fixed cell type, whereas current methods (`Citrus`, `CellCnn`, `cydar`, and `diffcyt`) learn cell types and perform differential analysis jointly. The narrower field of application allows us to define a more specific statistical model with easier to control statistical guarantees. Only the `Bioconductor workflow` by Nowicka et al. (2017) focuses on specific cell types, but as mentioned before, they employ marginal regression which makes comparison to our multiple regression method difficult—as the two methods have different aims.

We present two versions of multiple regression: (i) A Generalized Linear Model (GLM) for unpaired samples. A GLM is a regression model that allows for a response and error terms that follow different distributions than the normal. (ii) A restricted Generalized Linear Mixed Model (GLMM), which is a GLM that allows for random and fixed effects, for paired samples—when the same donor provides two samples, one for each condition. GLMs and GLMMs are generalizations of least squares to data from the exponential family. In our case, we will use logistic regression to model the experimental condition with a Bernoulli distribution and link it to a linear model of marker expressions with the logit function.

Our models depart from the classic model where the marker expressions are the response variables. In our GLMs, the experimental condition *Y* is independent of the *j*th marker expression *X*_*j*_ given the other markers *X*_−*j*_ (all makers *X*_1_, …, *X*_*P*_ except the marker *X*_*j*_) if and only if the *j*th regression coefficient is zero (for a mathematical proof of this statement see Proposition 2.2 in Candès et al. (2018)). In contrast, the usual marginal regression analysis does not allow for such conditional statements. For instance, it would not allow us to rule out markers that are merely correlated with other makers but are independent of the experimental condition—as illustrated with the example earlier.

In summary, our two main contributions are:

1. We present a conditional differential analysis to avoid biases arising from marker correlations by using multiple regression instead of marginal regression.
2. We present two multiple regression strategies that work with the unsummarized expression data to maximize statistical power and account for patient heterogeneity to safeguard against false discoveries: (i) GLMs with a patient-level bootstrap, and (ii) GLMMs with a patient-level random effect.

In Section 2, we review the statistical background for GLMs and GLMMs. Section 3 evaluates the statistical properties of both strategies implemented in our *R* package `CytoGLMM` on different simulated datasets, and illustrates the full workflow for real pregnancy data. In Section 4, we discuss our results in terms of biases and confounders.

## 2 Methods

### 2.1 Preprocessing

We recommend that marker expressions be corrected for batch effects (Nowicka et al. 2017; Chevrier et al. 2018; Schuyler et al. 2019; Van Gassen et al. 2020; Trussart et al. 2020) and transformed using variance stabilizing transformations to account for heteroskedasticity, for instance with a hyperbolic sine transformation with the cofactor set to 150 for flow cytometry, and 5 for mass cytometry (Bendall et al. 2011). This transformation assumes a two-component model for the measurement error (Rocke and Lorenzato 1995; Huber et al. 2003) where small counts are less noisy than large counts. Intuitively, this corresponds to a noise model with additive and multiplicative noise depending on the magnitude of the marker expression; see (Holmes and Huber 2019) for details.

### 2.2 Generalized Linear Model (GLM)

The goal of the GLM is to find protein expression patterns that are associated with the condition of interest, such as a response to a stimulus. We set up the GLM to predict the experimental condition from protein marker expressions, thus our experimental conditions are response variables and marker expressions are explanatory variables. The response variable *Y*_*i*_ is a binary variable encoding experimental condition as zero or one. The response variable can be modeled as a Bernoulli random variable with probability *π*_*i*_ for each cell. Then we use the logit link to relate the linear model to binary responses. The linear model predicts the logarithm of the odds of the *i*th cell being *Y*_*i*_ = 1 instead of *Y*_*i*_ = 0. The linear model has one coefficient per protein marker *β*_1_, …, *β*_*P*_ and an intercept *β*_0_. If *π*_*i*_ is 0.5 then the cell could have come from either *Y*_*i*_ = 1 or *Y*_*i*_ = 0 with equal probability. If *π*_*i*_ is either very close to one or zero, then the cell is strongly representative of a cell observed under *Y*_*i*_ = 1 or *Y*_*i*_ = 0, respectively. We observe the protein marker expressions ***x***_*i*_. For each cell we measure *P* protein markers.

The response probabilities *π*_*i*_ are not observed directly, only *Y*_*i*_ = *Y*_*i*_ and ***x***_*i*_ are observed. Note that ***x***_*i*_ is observed with errors. Here, we make the approximating assumption that the covariates are fixed. Our results will show that this assumption is conservative and introduces a regularization of the estimated coefficients. We estimate *π*_*i*_ from the data using maximum likelihood with the function `glm` in *R*. Our logistic regression model, which is part of a general class of GLMs, can be summarized in the following form:

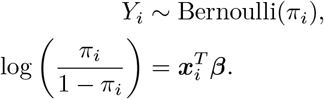

For likelihood inference, we use the nonparametric bootstrap and resample entire donors with replacement to preserve the cluster structure. At the cell-level, we resample cells with replacement within each donor. We build percentile confidence intervals and compute *p*-values by inverting the intervals assuming two-sided intervals with equal tails (Efron and Tibshirani 1994). We use the Benjamini-Hochberg (BH) (Benjamini and Hochberg 1995) and Benjamini–Yekutieli (BY) (Benjamini and Yekutieli 2001) procedures to control the False Discovery Rate (FDR). We refer to GLM with BH control as GLM-BH, and with BY control as GLM-BY.

### 2.3 Generalized Linear Mixed Model (GLMM)

We make additional modeling assumptions by adding a random effect term in the standard logistic regression model to account for the subject effect. The covariates ***x***_*ij*_ are the same as in the fixed effects GLM, except now we have an additional index *j* that indicates from which donor the cell was taken. Each cell *i* maps to a donor *j*. The additional term ***u***_*j*_ represents regression coefficients that vary by donor. The statistical model can be summarized as,

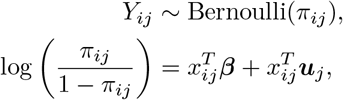

with a multivariate normal distribution and covariance matrix **Σ** for the random effect term ***u***_*j*_,

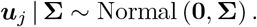

Analog to our GLM, we make the approximating assumption that the covariates are fixed.

The mixed effect model is a compromise between two extremes. On the one hand, we could estimate separate regression coefficients for each donor. This corresponds to random effects modeled with a multivariate normal distribution with infinite standard deviations and no constraint on how coefficients are related to each other. On the other hand, we could pool all donors into one group and ignore the donor information. This corresponds to a GLM with no random effects, with no additional variability besides the fixed effect term. A compromise between these two extremes is to estimate the standard deviations of the random effects from data, allowing the regression model to learn from the other donors. Mixed effects procedures are related to empirical Bayes procedures (Weber et al. 2019). The first s tep of an empirical Bayes procedure would estimate the mean and covariance matrix of the random effect term. The second step would fix the random effect parameters at their estimated values and estimate the fixed effect parameters. In contrast, t he mixed effect procedure estimates the parameters of both steps jointly. This is possible for flow and mass cytometry data because of the relatively small number of proteins.

We use the method of moments as implemented in the *R* package `mbest` to estimate the model parameters ***β***, ***u***_*j*_, and **Σ**. For likelihood inference, we use the asymptotic theory derived by Perry (2017). The author showed that the sampling distribution of the estimated parameters can be approximated by a normal distribution. We use this mathematical alternative to the bootstrap method to create approximate confidence intervals and *p*-values. As in the GLM case, we use the Benjamini-Hochberg (BH) and Benjamini–Yekutieli (BY) procedures to control the FDR. We refer to GLMM with BH control as GLMM-BH, and with BY control as GLMM-BY.

## 3 Results

We first evaluate these procedures for both paired and unpaired samples on simulated datasets. We then test them on a real pregnancy dataset.

### 3.1 Simulated Datasets

We generate simulated data with both cell and donor level variability. We allow for negative and positive correlations between markers and a wide range of correlation strengths. We simulate different scenarios ranging from weak to strong patient/cell variability. To make sure that we generate positive counts we use a Poisson noise model after transforming the generated expressions to positive real numbers using the exponential function. This is similar to using the log link function for Poisson GLMs. Overall, there are four main parameters: correlation *ρ*_*B*_ and standard deviation *σ*_*B*_ at the cell level, and correlation *ρ_U_* and standard deviation *σ*_*U*_ at the donor level. Additionally, we can regulate the number of cells per sample and the number of donors per dataset. The differential expression signal is induced by shifting the mean vector on the logarithmic scale.

We study the differential expression of three out of 10 markers after simulating exposure of cells to an experimental condition with two levels: stimulated versus unstimulated cells. We consider one underlying data generating mechanisms described by a hierarchical model:

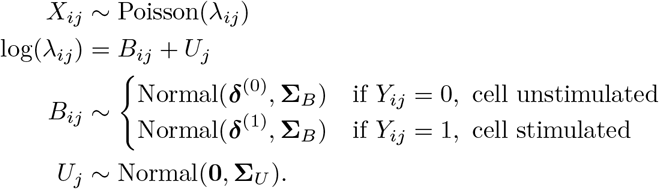

The stimulus activates three proteins and induces a difference in marker expression. We define the effect size to be the difference between expected expression levels of stimulated versus unstimulated cells on the log-scale. We choose a set *C* = 3 of active markers. All markers that belong to the active set, have a non-zero effect size, whereas, all markers that are not, have a zero effect size:

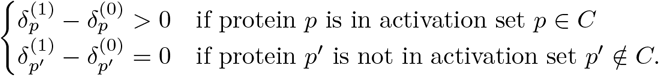

Both covariance matrices have an autoregressive structure,

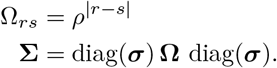

We regulate two separate correlation parameters: a cell-level *ρ*_*B*_ and a donor-level *ρ*_*U*_ coefficient. Non-zero *ρ*_*B*_ or *ρ*_*U*_ induce a correlation between condition and marker expression even for markers with a zero effect size.

We performed simulations with a variety of different parameters. All simulations have 16 samples. For paired samples, those 16 samples come from 8 donors. For unpaired samples, those 16 samples come from 16 donors. Each sample has 1,000 cells. We compared the observed FDR and the power. The FDR measures the statistical type 1 errors, the expected proportion of falsely declared discoveries over the total number of reported discoveries. The statistical power represents the proportion of correctly reported discoveries over the total number of true discoveries.

Figures 1 and 2 show a summary averaged over 100 runs for paired sample and unpaired sample experiments with effect size 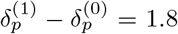 and 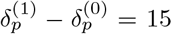, respectively, and varying standard deviation *σ* and correlation *ρ* parameters. The dashed lines indicate the target FDR of 0.05.

**Figure 1:**
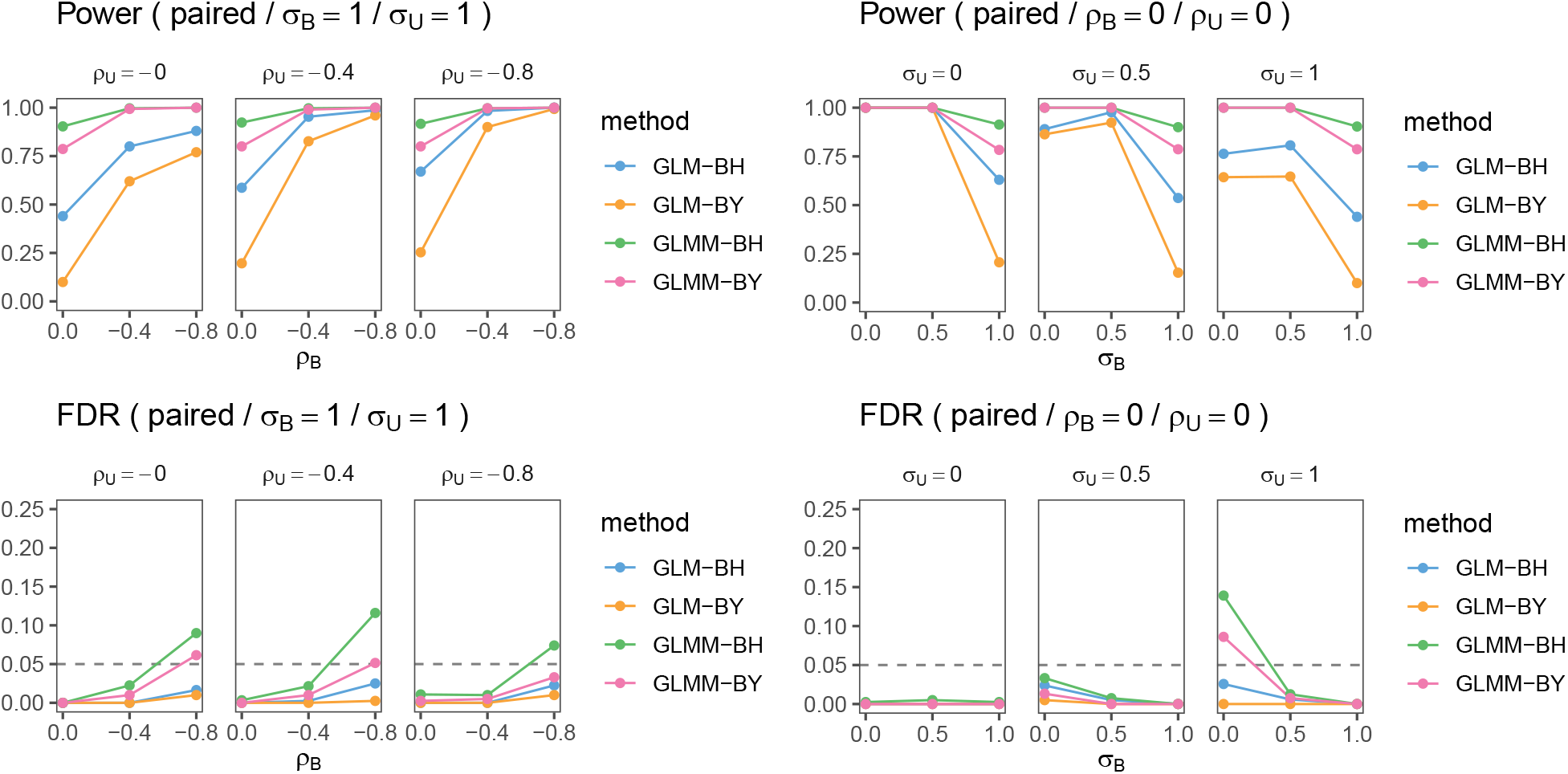
Summary of experiments with 1,000 cells per sample averaged over 100 runs. The horizontal dashed line represents the target FDR. Postfixes BH and BY stand for the respective FDR control procedure. Subscripts *B* and *U* indicate cell and donor-level standard deviation *σ* and correlation *ρ*, respectively.

**Figure 2:**
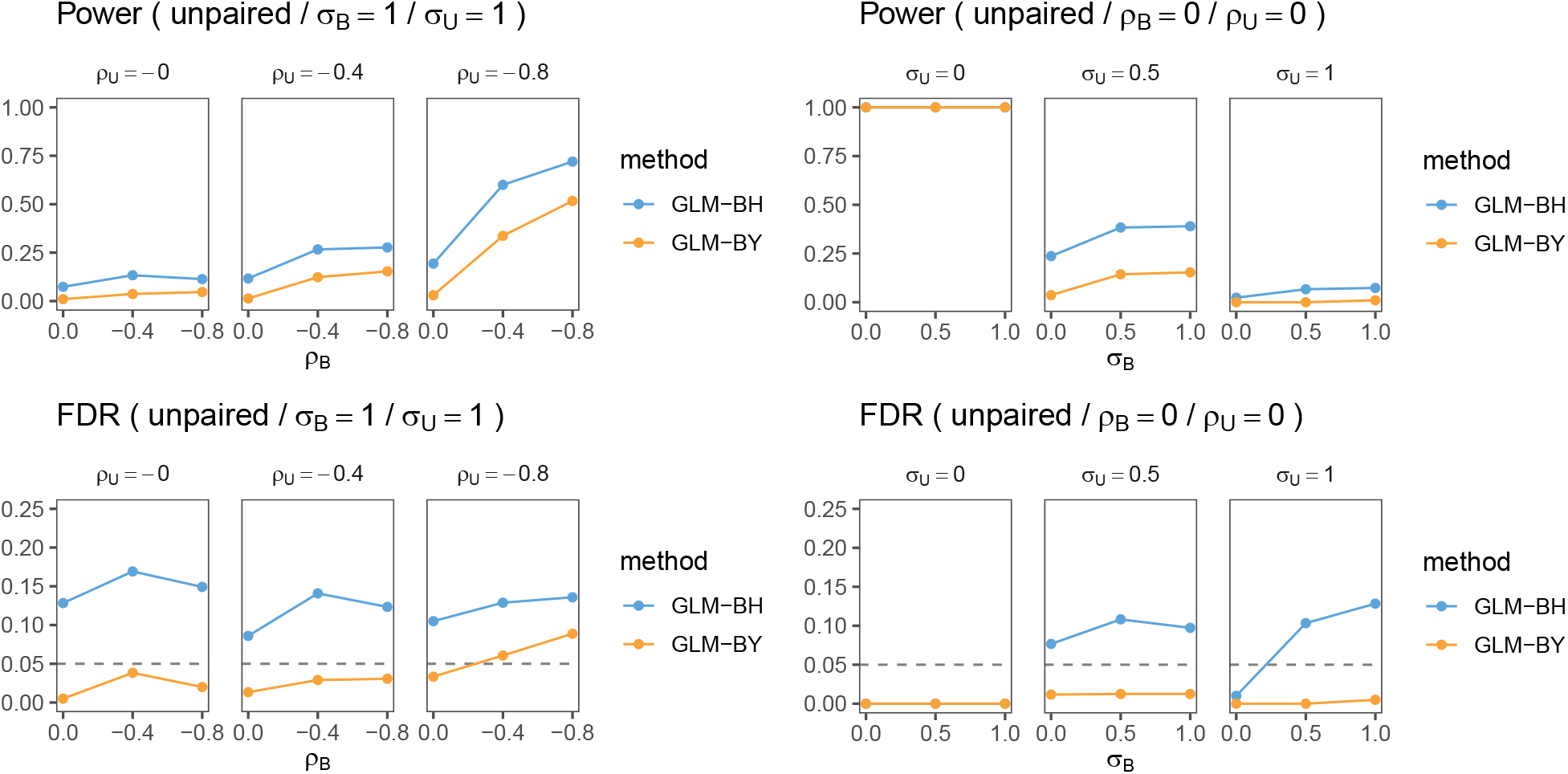
Summary of experiments with 1,000 cells per sample averaged over 100 runs. The horizontal dashed line represents the target FDR. Postfixes BH and BY stand for the respective FDR control procedure. Subscripts *B* and *U* indicate cell and donor-level standard deviation *σ* and correlation *ρ*, respectively.

First, let’s consider the paired samples experiment. The plots on the left show results when we vary cell and donor-level correlations at a fixed amount of cell *σ*_*B*_ = 1 and donor *σ*_*U*_ = 1 marker standard deviations. We observe only small differences across donor correlations *ρ*_*U*_ of a small increase of power with increasing correlation. In contrast, there are large increases of power as a function of cell correlations *ρ*_*B*_. In the panel of plots on the right, we set both correlations to zero and vary the marker standard deviations. In this setting, we again observe major changes with increasing standard deviations at the cell-level *σ*_*B*_: the larger the cell-level variability, the lower the power. This is also true for donor-level variability, though to a much lesser extent. FDR is controlled below its target level under medium cell-level marker correlations (|*ρ*_*B*_| ≤ 0.4) except when cell variability is at zero *σ*_*B*_ = 0, and donor variability is at one *σ*_*U*_ = 1. As expected, the BY procedure is more conservative than the BH procedure, that is both FDR and power are lower. Interestingly, power increases with cell-level correlations *ρ*_*B*_, and is virtually unaffected by donor-level correlations *ρ*_*U*_. Overall, GLMM methods are more powerful than GLM methods. Figure 3 shows simulations for power and FDR with varying numbers of paired samples. Both cell and donor standard deviations are set to *σ*_*B*_ = *σ*_*U*_ = 1, and correlations are set to *ρ*_*B*_ = *ρ*_*U*_ = 0. An efficiency gain is clearly visible when we compare how many paired samples are needed to achieve 80% power. We observe that for GLMMs we need 7 paired samples to exceed the 80% power threshold, whereas for GLMs we need 13 paired samples to achieve the same.

**Figure 3:**
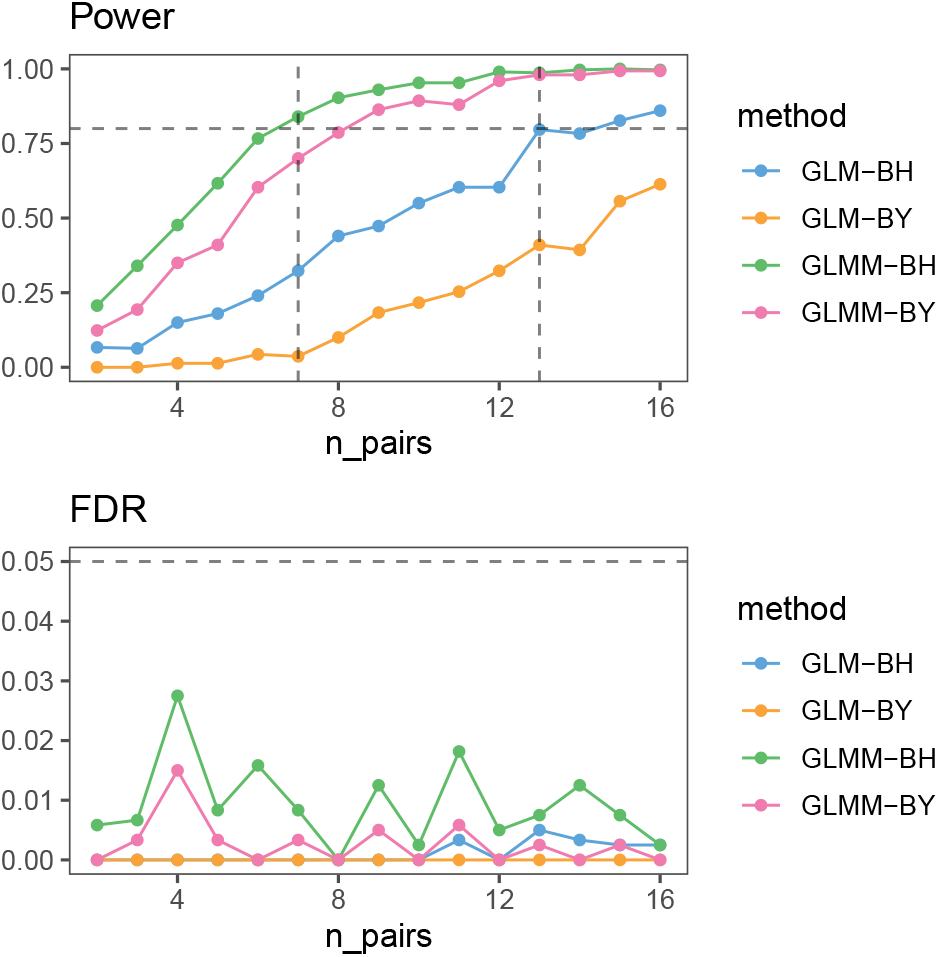
Summary of experiments with 1,000 cells per sample averaged over 100 runs. Power: The horizontal dashed line represents a power of 0.8. FDR: The horizontal dashed line represents the target FDR of 0.05.

In the unpaired samples experiment, we only show GLM results as the GLMM results have zero power, there is no data to estimate the donor-level random effect term. We observe up to 20% FDR with a target FDR of 5%. To have non-zero power we need to increase the effect size to 15 (in comparison, for paired experiments the effect size is set to 1.8). Furthermore, FDR is only controlled under medium cell-level marker correlations using the more conservative BY procedure, with BH exceeding 0.05 in most scenarios except when we have zero donor-level variability *σ*_*U*_ = 0. As before, BY comes with a loss of power.

### 3.2 Experimental Dataset

We reanalyze a recently published dataset on the maternal immune system during pregnancy (Aghaeepour et al. 2017). The study provides a rich mass cytometry dataset collected at four time points during pregnancy in two cohorts. The authors isolated cells from blood samples and stimulated them with several activation factors. The goal was to explain how immune cells react to these stimuli, and how these reactions change throughout pregnancy. Findings from such experiments might identify immunological deviations implicated in pregnancy-related pathologies.

The authors collected data at early, mid, late pregnancy, and six weeks postpartum. Samples were left unstimulated or stimulated. Stimulation conditions included: interferon-*α*2A (IFN*α*), lipopolysaccharide, and a cocktail of interleukins (ILs) containing IL-2 and IL-6. They processed the samples on a CyTOF 2.0 mass cytometer instrument, and bead normalized the data to account for signal variation over time from changes in instrument performance (Finck et al. 2013).

In our analysis, we focus on comparing early (first trimester, *Y*_*i*_ = 0) with late (third trimester, *Y*_*i*_ = 1) pregnancy samples stimulated with IFN*α* in the first cohort of 16 women. We gate cells into cell types and organize them in a data frame. We follow the gating scheme detailed in (Aghaeepour et al. 2017) and define 12 cell types using the *R* package `openCyto` (Finak et al. 2014): memory CD4 positive T cells (CD4+Tmem), naive CD4 positive T cells (CD4+Tnaive), memory CD8 positive T cells (CD8+Tmem), naive CD8 positive T cells (CD8+Tnaive), *γδ*T cells (gdT), regulatory T memory cells (Tregsmem), regulatory T naive cells (Tregsnaive), B cells, classical monocytes (cMC), intermediate monocytes (intMC), non-classical monocytes (ncMC), and Natural Killer cells (NK). Out of the 32 protein markers measured on each cell, the authors defined 22 markers as gating markers, and 10 as functional markers. The functional markers are pSTAT1, pSTAT3, pSTAT5, pNF*κ*B, total I*κ*B, pMAPKAPK2, pP38, prpS6, pERK1/2, and pCREB (in plots Greek symbols are replaced by Latin symbols).

We plot the maximum likelihood (GLM) and the method of moments estimates (GLMM) with 95% confidence intervals for the fixed effects ***β*** (Figure 4). The estimates are on the log-odds scale. We see that pSTAT1 is a strong predictor of the third trimester. This means that one unit increase in the transformed marker expression makes it between exp(1) = 2.7 to exp(1.5) = 4.5 (95% confidence interval for GLMM) more likely to be a cell from the third trimester, while holding the other markers constant. pSTAT3 and pSTAT5 have negative coefficients. This means pSTAT3 and pSTAT5 predict the first trimester, while holding the other markers constant. Only pSTAT1, pSTAT3, and pSTAT5 are below an FDR of 0.05. Our results corroborate previous findings by Aghaeepour et al. (2017) reporting an increase of pSTAT1 during the third trimester for IFN*α* stimulated samples.

**Figure 4:**
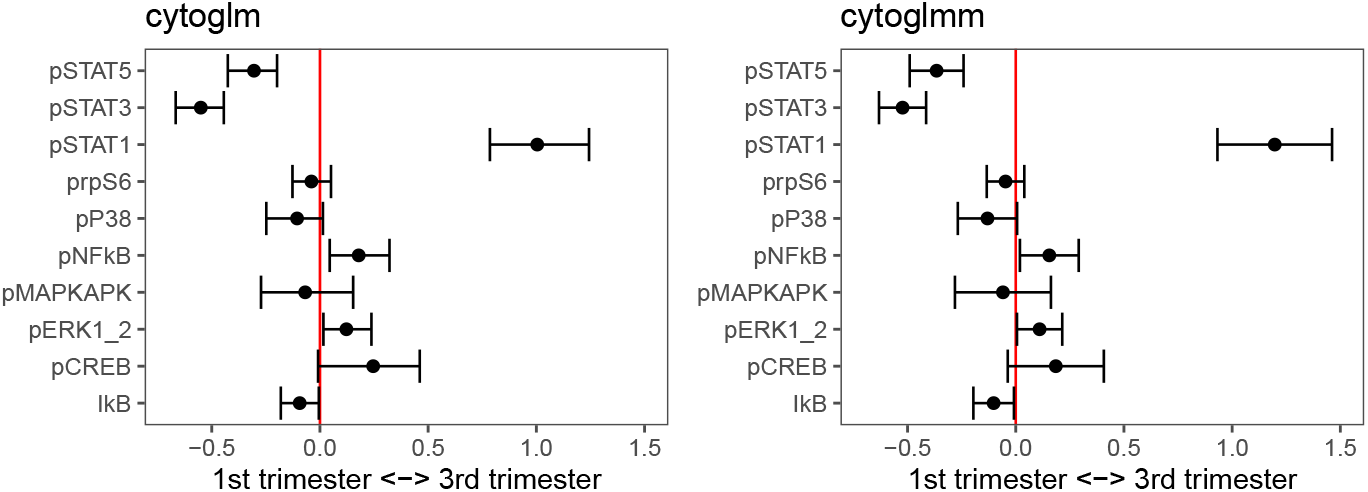
Methods comparison between bootstrap GLM (cytoglm) and GLMM (cytoglmm). The horizontal axes are on the log-odds scale. The vertical axes are the protein markers.

## 4 Discussion

Our new *R* package `CytoGLMM` is applicable to a wide range of cytometry studies. Besides comparisons on paired samples, where samples are available for the same subject under different experimental conditions, our `CytoGLMM` is applicable to unpaired samples, where samples are collected on two separate groups of individuals.

Our simulation experiments compare multiple regression GLM and GLMM, as implemented in cytoglm and `CytoGLMM` in our *R* package. In simulated paired samples experiments, both GLMM-BH and GLMM-BY procedures control the FDR below the target FDR under cell-level marker correlations with an autoregressive structure with correlations up to ±0.4. GLMM methods are more powerful than GLM methods for paired samples. GLMM methods can account for the patient-to-patient variation in the model, whereas GLM methods treat this variation as noise, which results in noisier and thus less powerful estimates. For unpaired samples, we are forced to use the nonparametric bootstrap method for GLMs because there are no paired samples available to estimate the random effect term. In simulated unpaired experiments, only BY controls the target FDR. In practice, this means that we need a much higher donor samples size to detect a differential expression compared to paired experiments.

Overall, larger cell-level and donor-level correlations increase power and reduce the observed FDR. Hypothesis testing under arbitrary dependency structure is still an active research topic (Barber, Candès, and others 2015; Candès et al. 2018; Fithian and Lei 2020). What is easier to explain is the reduction in power and FDR as a function of increased cell-level variance. Research in measurement error models shows that increased uncertainty in measured covariates is linked to biased estimates. The coefficient estimates are regularized—shrinking them towards zero—which translates into more conservative *p*-values; for extensive literature on this topic see Fuller (1987) and Carroll et al. (2006). In GLMMs, donor-level correlations have only a weak impact on power and observed FDR because we explicitly model correlations with a random effect term.

In general, biases in coefficient estimates of GLMs and GLMMs can occur when we leave out proteins from the analysis. Assume that we would like to relate variable protein *X* to experimental condition *Y*. If there exists a second protein *Z* both related to *X* and *Y*, then *Z* is called a confounder, and not including it in the analysis can change the coefficients estimates. In the pregnancy data, if we removed pSTAT1 from the analysis, the confidence intervals of pSTAT3 and pSTAT5 could change. Such a difference is expected if pSTAT1 is a confounder. If pSTAT1 is not a confounder, the coefficient estimates for pSTAT3 and pSTAT5 will be the same whether pSTAT1 is included or not. The change of coefficients depending on what markers are included in the model can have strong effects. We have observed that in some real datasets that one marker can make other markers change their sign depending on whether we include them or not. In such cases, we recommend keeping all markers in the analysis to avoid introducing confounding biases.

Our simulations are limited to a Poisson mixed effect model for protein marker expression. Our conclusions are only valid with respect to this model. The real data generating process might be different. Two main caveats are to be noted. First, we can only encode an experimental design comparing two groups. Second, we require gated cell types. To reduce the person-to-person bias of manual gating, we employed the *R* package `openCyto` (Finak et al. 2014). The curse of dimensionality makes it challenging to scale this approach to very high dimensional gating schemes.

A possible alternative to GLMMs are Generalized Estimating Equations (GEEs). GEEs are statistically more efficient when the covariance structure of the residuals are known. In our case, the covariance structure is unknown and needs to be estimated from the data. In most immunology studies, we only have a few donors without a given covariance structure (e.g. no time dependency), resulting in a hard and possibly unstable covariance estimation problem, which could result in an overall loss of efficiency (Wakefield 2013).

We applied `CytoGLMM` in wide range of immunology studies: comparison between influenza strains (Kronstad et al. 2018), comparison between pregnant and non-pregnant women (Le Gars et al. 2019), comparison between healthy controls and HIV+ individuals (Vendrame et al. 2020), comparison between multiple sclerosis patients treated with daclizumab beta or placebo (Ranganath et al. 2020), and comparison between Beninese sex workers and healthy controls (Zhao et al. 2020). Our next step is extending `CytoGLMM` to include more complicated experimental designs; e.g. twin studies (Brodin et al. 2015).

## Acknowledgments

This work was supported by the National Institutes of Health [U01AI131302 to C.A.B. and S.H., R56AI124788 to C.A.B. and S.H., R21AI130523 to C.A.B. and S.H., DP1DA046089 to C.A.B., R21AI130532 to C.A.B., R01AI133698 to C.A.B., R21AI135287 to C.A.B., 5T32AI007290-29 to L.M.K., TL1TR001084 to E.V., T32AI007502 to E.V., 1F32AI126674 to L.J.S.]; an A.P. Giannini fellowship [to L.M.K.]; and a Stanford Child Health Research Institute postdoctoral fellowship [to M.L.G.]. C.A.B. is the Tashia and John Morgridge Endowed Faculty Scholar in Pediatric Translational Medicine from the Maternal Child Health Research Institute, and a Chan Zuckerberg Investigator.

## Supplementary Material

### CytoGLMM R Package

Our *R* package is available on GitHub:

- https://github.com/christofseiler/CytoGLMM/

A vignette is available on our *R* package website:

- https://christofseiler.github.io/CytoGLMM/

### CytoGLMM Workflow

Here we illustrate a complete `CytoGLMM` workflow in *R*. Prepare data frame marker counts and sample information.

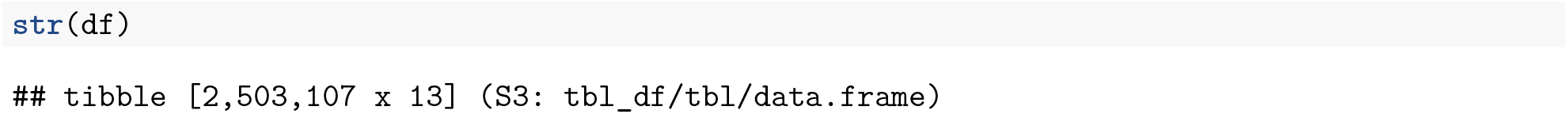
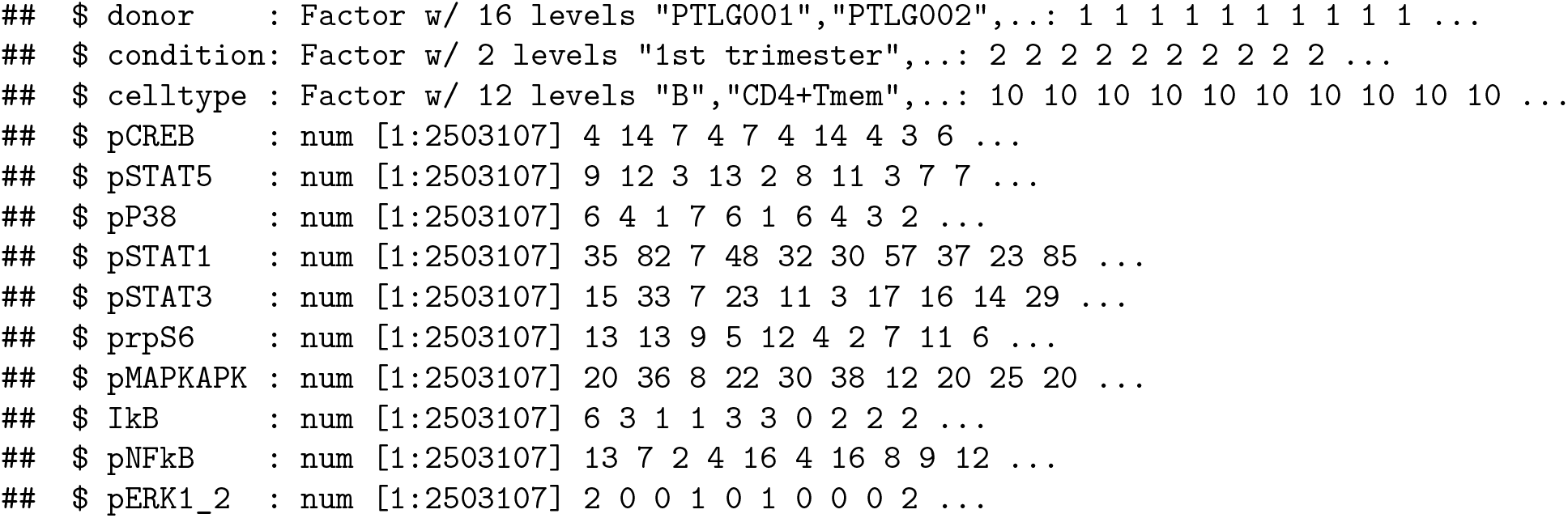

Select functional marker of interest.

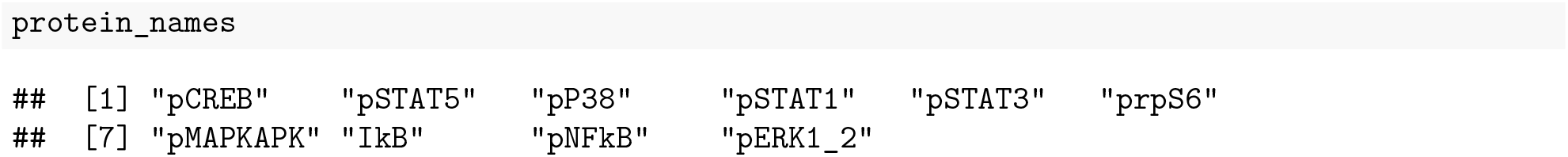

Transform counts.

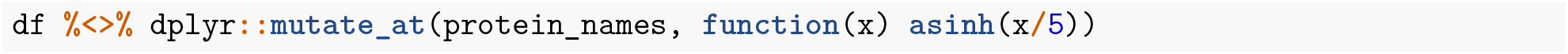

Subset to one celltype.

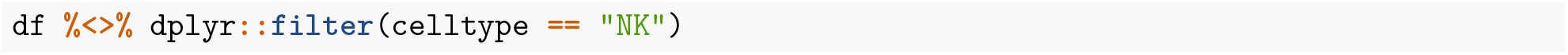

Fit the `cytoglm` model with 1000 bootstrap samples.

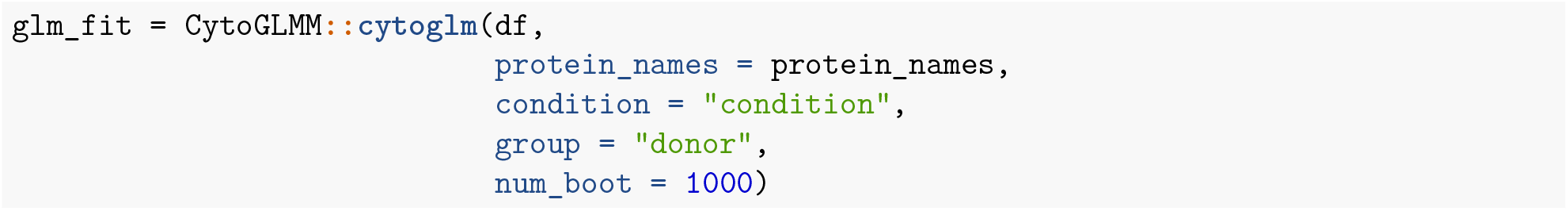

Fit the `cytoglmm` model.

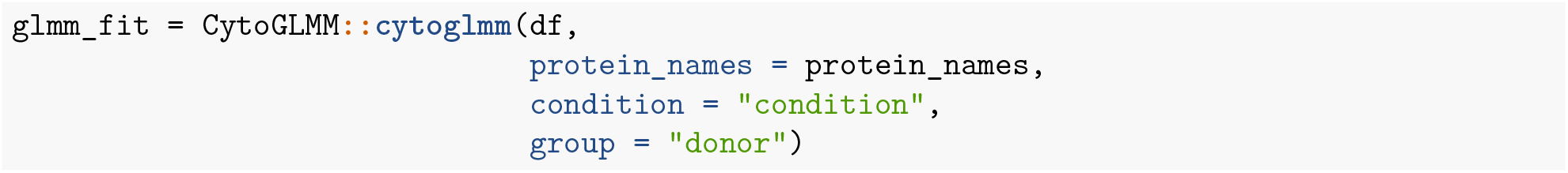

Use `print(glm_fit)` or `print(glmm_fit)` on the fitted object to obtain some additional details of the model that we just fitted.

Plot differential analysis results.

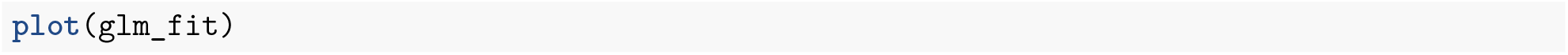

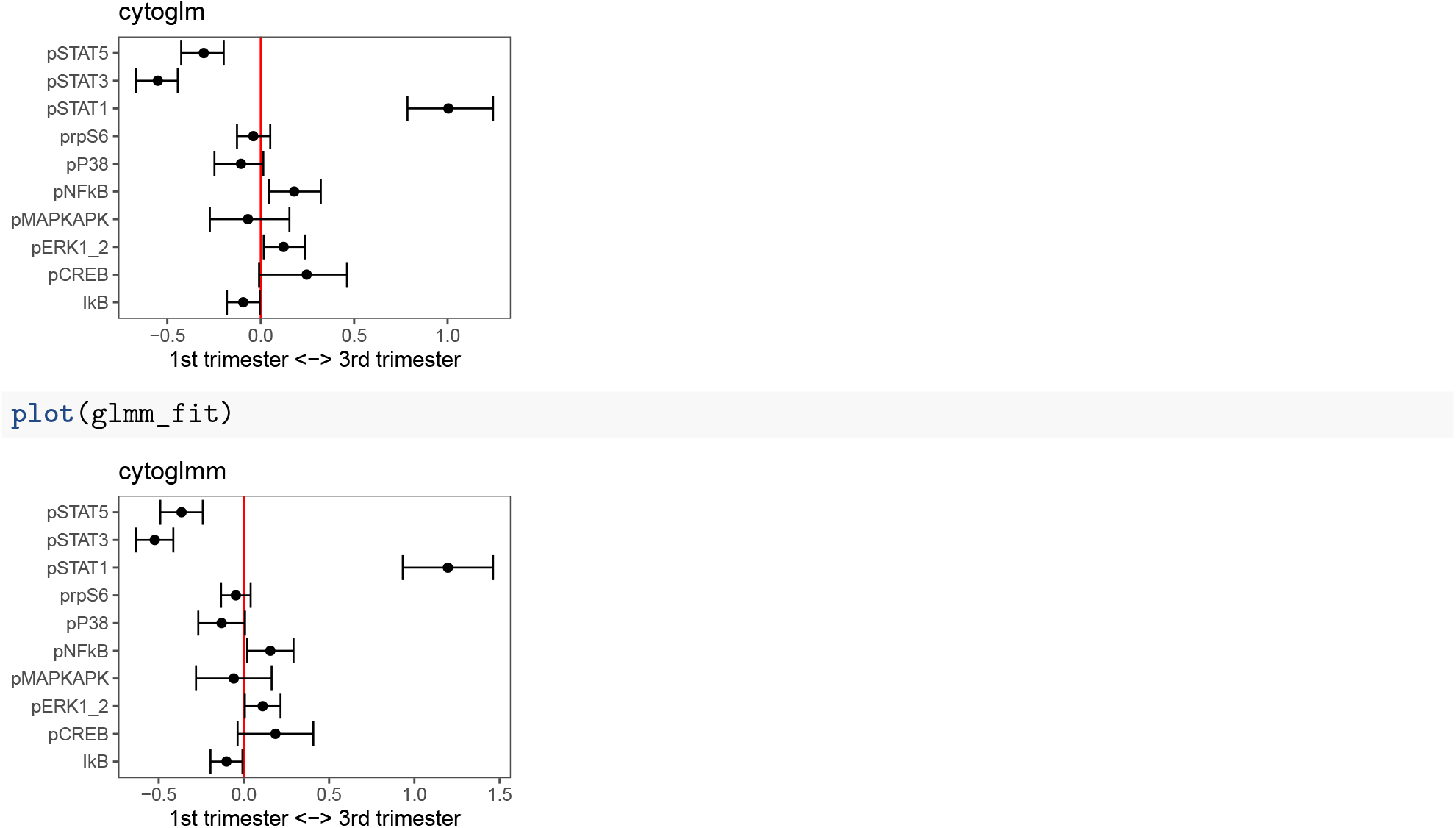

Extract *p*-values.

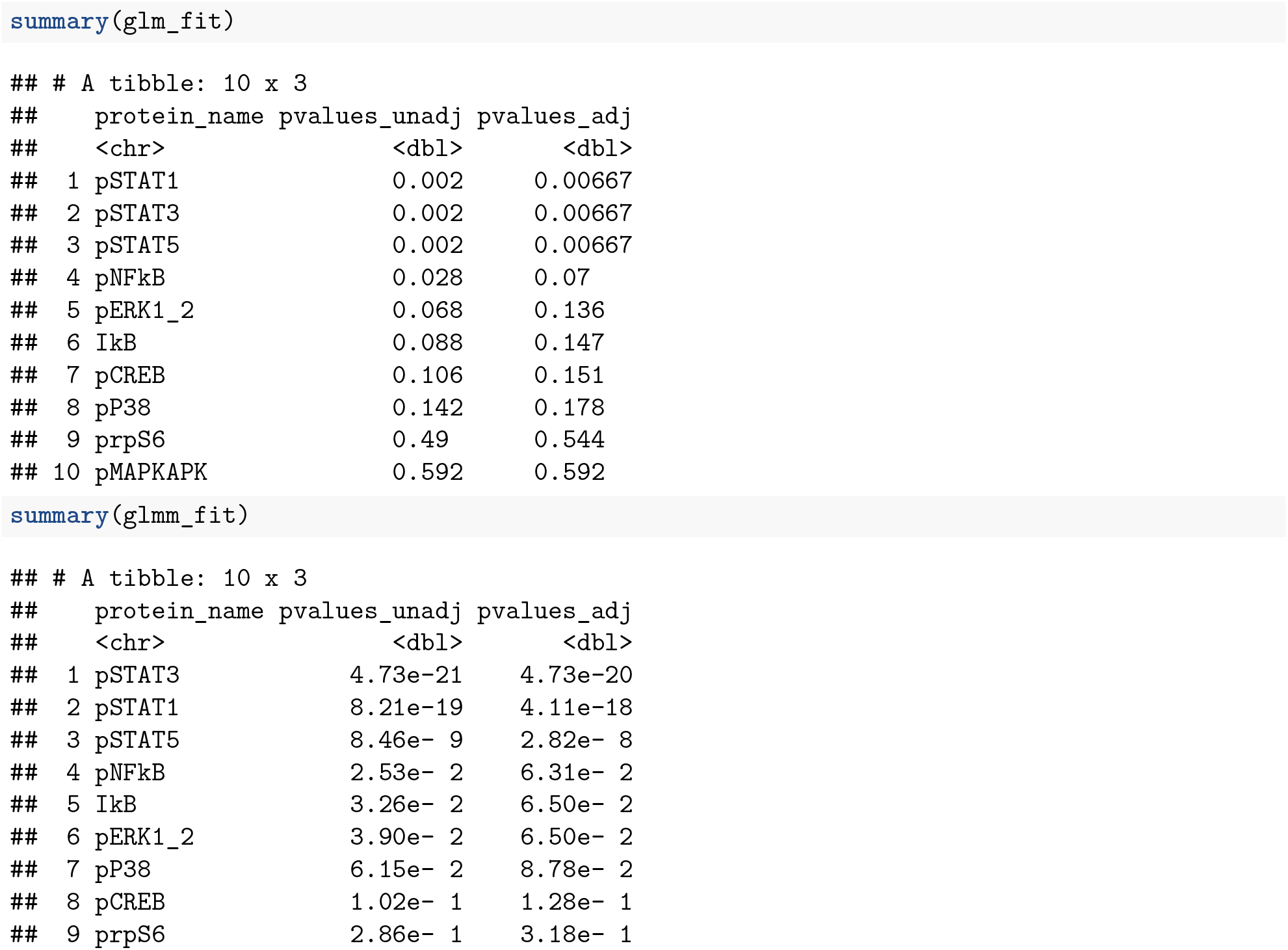
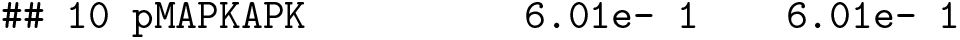

Filter adjusted p-values at some threshold.

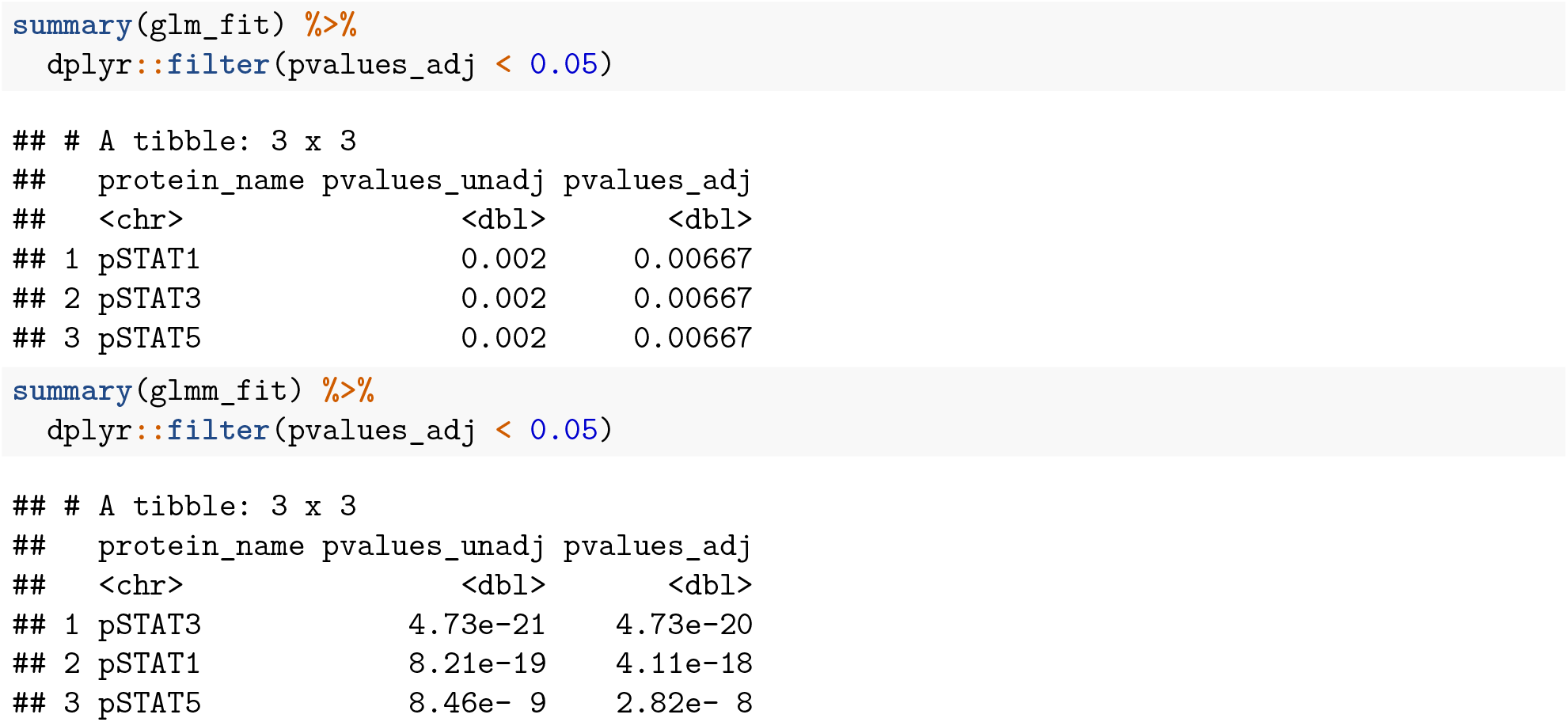

